# Plasticity in Na^+^/K^+^-ATPase thermal kinetics drives variation in the critical thermal minimum of adult *Drosophila melanogaster*

**DOI:** 10.1101/2022.08.31.506053

**Authors:** Mads Kuhlmann Andersen, R. Meldrum Robertson, Heath A. MacMillan

**Affiliations:** Department of Biology, Carleton University, Ottawa, ON, Canada; Department of Biology, Queen’s University, Kingston, ON, Canada

**Keywords:** Critical thermal minimum, sodium pump, enzyme kinetics, ionoregulation, glia

## Abstract

The majority of insects can acclimate to changes in their thermal environment and counteract temperature effects on neuromuscular function. At the critical thermal minimum a spreading depolarization (SD) event silences central neurons, but the temperature at which this event occurs can be altered through acclimation. SD is triggered by an inability to maintain ion homeostasis in the extracellular space in the brain and is characterized by a rapid surge in extracellular K^+^ concentration, implicating ion pump and channel function. Here, we focused on the role of the Na^+^/K^+^-ATPase specifically in lowering the SD temperature in cold-acclimated *D. melanogaster*. After first confirming cold acclimation altered SD onset, we investigated the dependency of the SD event on Na^+^/K^+^-ATPase activity by injecting an inhibitor, ouabain, into the head of the flies to induce SD over a range of temperatures. Latency to SD followed the pattern of a thermal performance curve, but cold acclimation resulted in a left-shift of the curve to an extent similar to its effect on the SD temperature. With Na^+^/K^+^-ATPase activity assays and immunoblots, we found that cold-acclimated flies have ion pumps that are less sensitive to temperature, but do not differ in their overall abundance in the brain. Combined, these findings suggest a key role for plasticity in Na^+^/K^+^-ATPase thermal sensitivity in maintaining central nervous system function in the cold, and more broadly highlight that a single ion pump can be an important determinant of whether insects can respond to their environment to remain active at low temperatures.

## Introduction

Temperature directly influences insect performance; as temperature is lowered most biological processes are slowed (Lee, 2012). The majority of insects are chill susceptible, meaning that adverse effects of cold on physiology begin to manifest at temperatures above those that cause any freezing (Bale, 1996; Sinclair, 1999; Overgaard and MacMillan, 2017). When cooled to their critical thermal minimum (CT_min_), these insects lose the ability to perform coordinated movements and shortly after enter a paralytic state known as chill coma (Mellanby, 1939; Hazell and Bale, 2011; MacMillan and Sinclair, 2011; Overgaard and MacMillan, 2017). While in chill coma, an insect’s potential fitness is effectively zero, and it is therefore not surprising that this temperature threshold closely relates to geographical distribution (Addo-Bediako et al., 2000; Bale, 2002; Sunday et al., 2011). Within the genus *Drosophila*, for example, chill coma temperatures vary markedly among species and closely correlate with poleward limits to species distribution (Kimura, 2004; Kellermann et al., 2012; Andersen et al., 2015b).

The current mechanistic model for chill coma onset posits that a cold-induced spreading depolarization (SD) event shuts down nervous system function, causing the loss of coordinated movements observed at the CT_min_ (Robertson et al., 2017; Andersen et al., 2018). Subsequently, low temperature decreases muscle excitability, which ultimately leads to complete paralysis and chill coma onset (Hosler et al., 2000; MacMillan et al., 2014; Findsen et al., 2016). However, not all insect species experience this loss of muscle excitability while still entering chill coma (Andersen et al., 2015a), and as such only experience a relatively “shallow” coma caused by the SD event in the central nervous system (Andersen and Overgaard, 2019). Similar observations have been made for insect comas induced by heat (Money et al., 2005; Jørgensen et al., 2020) and anoxia (Rodgers et al., 2007), implying SD is a common consequence of abiotic stress in insects.

Cold-induced SD and chill coma are plastic traits that can vary greatly. This is also true for *Drosophila*, where the chill coma and SD temperatures cover a range of more than 10°C among species (Mellanby, 1954; Kellermann et al., 2012; Andersen et al., 2015b; MacMillan et al., 2015b) and by several degrees within a species depending on thermal history (Kelty and Lee Jr, 1999; Overgaard et al., 2011; Ransberry et al., 2011; Armstrong et al., 2012; MacMillan et al., 2015a; Schou et al., 2017; Andersen et al., 2018). This makes drosophilids, such as *Drosophila melanogaster*, excellent model systems for studying the physiological mechanisms underlying variation in cold tolerance. Indeed, research conducted over the last decade has relied heavily on this clade to make great progress in describing the processes that underlie improved organismal cold tolerance (see reviews by Overgaard and MacMillan (2017) and Overgaard et al. (2021)).

While SD is a well-described phenomenon, the specific mechanism underlying the SD event itself remains unknown despite nearly eight decades of research in mammalian model systems (Leao, 1944) and two decades in insects (Robertson, 2004). A central feature of SD is a rapid disruption of extracellular ion homeostasis in the interstitium of the brain (Hansen and Zeuthen, 1981; Robertson, 2004; Rodgers et al., 2010). This disruption is characterized by a surge in extracellular K^+^ concentration associated with a near-complete depolarization of affected neurons and glia that effectively silences the area of the central nervous system experiencing the SD (Pietrobon and Moskowitz, 2014; Robertson et al., 2020). Thus, mechanisms responsible for extracellular ion homeostasis maintenance in the brain appear to be at least partially involved in the SD mechanism. While SD can be induced by abiotic stress, it can also be induced by a range of pharmacological interventions, common to which is that they either directly or indirectly inhibit activity of Na^+^/K^+^-ATPase, a key regulator of ion balance regulation in the central nervous system (Treherne and Schofield, 1981; Rodgers et al., 2010; Spong et al., 2016a; Andrew et al., 2022). Thus, the Na^+^/K^+^-ATPase appears to play a central role in the dynamics of SD induction, and by extension that the ability to maintain Na^+^/K^+^-ATPase activity when homeostasis is challenged by cold exposure could mitigate or delay the onset of SD (Rodgers et al., 2010; Spong and Robertson, 2013). Indeed, the findings of Cheslock et al. (2021) suggest that an acclimation treatment that lowers the SD temperature also lowers the thermal sensitivity of Na^+^/K^+^-ATPase activity. Thus, the ability to maintain ion homeostasis, both locally and systemically, appears critical for insect cold tolerance (Overgaard and MacMillan, 2017; Overgaard et al., 2021).

In the present study we investigate the proposed link between improved cold tolerance (i.e. lower cold-induced SD temperature) and thermal sensitivity of brain Na^+^/K^+^-ATPase. To do so, we induced phenotypic plasticity in the cold tolerance in *Drosophila melanogaster* by acclimating them to either 15°C or 25°C throughout development. Once variation in cold tolerance was confirmed, we investigated how temperature affected the Na^+^/K^+^-ATPase in *D. melanogaster* brains and how this might relate to the SD temperature. Here, we hypothesized that improved cold tolerance of the cold-acclimated group would be enabled by a lowered thermal sensitivity of Na^+^/K^+^-ATPase in the brain, such that neural function is better maintained at critically low temperatures. We further hypothesized that cold tolerance plasticity is facilitated by a greater abundance of Na^+^/K^+^-ATPase protein itself in the brain. We investigate this by 1) estimating how effectively Na^+^/K^+^-ATPase assisted in preventing SD *in vivo* across a range of temperatures, 2) measuring the activity of Na^+^/K^+^-ATPase in an enzyme-linked assay *in vitro* down to SD-inducing temperatures, and 3) determining the relative abundance of Na^+^/K^+^-ATPase protein in brains collected from cold- and warm-acclimated flies.

## Materials and Methods

### Animal husbandry

The line of *Drosophila melanogaster* (Meigen, 1830) used in this study was established from isofemales collected in London and Niagara on the Lake (Ontario, Canada) in 2007 (Marshall and Sinclair, 2010). Adult flies were reared in 200 mL bottles containing ∼ 40 mL banana-based medium (recipe: 0.95 L of water, 8 g agar, 27.5 g active yeast, 2 g methylparaben, 137.5 g organic banana, 47.5 g corn syrup, 30 g liquid malt, 3 mL propionic acid) and kept at 18°C with a 12h:12h light:dark cycle. Flies for experiments were produced by letting adult flies oviposit for 4-6 h in a new bottle with medium under rearing conditions, which resulted in rearing densities of 150-200 flies per bottle. The new egg-containing bottle was then transferred to a temperature-controlled incubator (MIR-154-PA, Panasonic) where the eggs were allowed to develop into adults at 15 or 25°C. Newly emerged adults were transferred to 40 mL vials containing ∼ 7 mL of the same medium. After maturing for 6-9 days at their respective acclimation temperatures, males were discarded and only females were used in experiments (to avoid potential sex-specific differences and because the larger size of females makes dissections easier). No anesthesia was used to separate females from males in the present study as all experiments involved individual flies being handled under a microscope or dissected, meaning that flies are free from potential effects of sorting under CO_2_ (Nilson et al., 2006; MacMillan et al., 2017).

### Preparation for electrophysiology

To prepare for electrophysiological experiments, flies were gently held by the head with a 100 μL pipette tip attached to an aspirator, immobilized in a thin layer of wax on a glass cover slide and further secured with a bent minuten pin gently positioned between the thorax and abdomen and fixed in the wax. After immobilization, micro scissors were used to cut a small hole in the head cuticle to access the brain, and a small incision was made in the abdomen to insert an Ag/AgCl wire for grounding (Spong et al., 2016b).

### Measurement of cold-induced spreading depolarization

In this study, we used the spreading depolarization (SD) temperature as an indirect measure of the whole-animal CT_min_, because this event has been found to be coincident with the CT_min_ in *Drosophila* (Andersen et al., 2018). The onset of SD was measured electrophysiologically with glass microelectrodes: Filamented glass capillaries (1B100F-4, World Precision Instruments (WPI), Sarasota, Florida, USA) were pulled on a Flaming-Brown P-1000 micropipette puller (Sutter Instruments, Novato, CA, USA) to a tip resistance of 5-7 MΩ when backfilled with 500 mM KCl. The filled glass microelectrodes were placed in electrode holders containing a Ag/AgCl wire, which were attached to micromanipulator (M3301-M3-L, WPI) and connected to a Duo 773 electrometer (WPI). Raw voltage outputs from the electrometer were digitized using a PowerLab 4SP A/D converter (ADInstruments, Colorado Springs, CO, USA) before being read by a computer running LabChart 4.0 software (ADInstruments).

For experiments, the glass cover slide with a fly attached was placed on a custom-built thermoelectrically cooled Peltier plate. Next, a type K thermocouple was placed immediately next to the fly’s head to monitor brain temperature (connected to the A/D converter with a thermocouple-to-analog converter (TAC80B-K; Omega, Stamford, CT, USA) and a glass microelectrode was inserted into the brain, through the hole cut in the head capsule, by gently advancing the electrode with the micromanipulator. Successful placement of the electrode was determined when the advancement of the microelectrode resulted in a small increase in the recorded voltage (5-10 mV); this was interpreted as having penetrated the glial blood-brain barrier such that the electrode was monitoring the transperineurial potential (Robertson et al., 2020; Cheslock et al., 2021). Subsequently, the temperature of the fly’s head was lowered from room temperature (22-23°C) by ∼ 1°C min^-1^ under manual control until an abrupt drop in the transperineurial potential was observed (30-50 mV in 5-10 s), indicative of SD having occurred. The temperature of SD onset was measured at the half-amplitude of the drop in transperineurial potential, and the amplitude of the event was calculated as the difference in potential before and after the onset. Furthermore, the descending slope at the half-amplitude of the drop in transperineurial potential was recorded (using the *Slope* function in the LabChart software). After SD onset, temperature was further lowered by 1°C after which flies were returned to room temperature, during which the transperineurial potential was also monitored to ensure that they could recover from the SD (i.e. the cold exposure wasn’t terminal). The SD temperature and amplitude were measured in ten flies from each of the 15°C and 25°C acclimation groups.

### Latency to spreading depolarization following ouabain injection

To determine whether SD onset characteristics were dependent on Na^+^/K^+^-ATPase activity, we injected ouabain into the head of the fly while continuously monitoring the transperineurial potential (as described above) (Rodgers et al., 2007; Rodgers et al., 2010; Spong et al., 2016b). Ouabain (O3125, Sigma, St. Louis, MO, USA) was diluted in saline (147 mM NaCl, 10 mM KCl 4 mM CaCl, and 10 mM HEPES at pH 7.2) to a concentration of 0.1 mM and slowly injected into the head of the fly in a small aliquot (∼ 14-15 nL) using a Nanoject II Auto-Nanoliter Injector (Drummond Scientific Company, Broomall, PA, USA). Following injection we measured the latency to the first SD event (there were often multiple events following the first one; example trace in **Fig. 2**). This measurement was performed at temperatures ranging from 10°C to 35°C in 5°C increments, and fly preparations were allowed to equilibrate temperature for 5-10 min before injection. We found that 10°C was too close to the SD temperature for the warm-acclimated group as it often led to a cold-rather than ouabain-induced SD, and 35°C was included for the warm-acclimated group so that the same number of temperatures were tested in each group. Six to eight flies from each acclimation group were measured at each temperature (N = 68 flies in total).

### Collection of brain samples

Flies were briefly sedated in 70% ethanol and subsequently dissected in standard *Drosophila* saline (in mM: 122.5 NaCl, 15 KCl, 2 CaCl_2_, 8.5 MgCl_2_, 10.2 NaHCO_3_, 4.3 NaH_2_PO_4_, 15 MOPS, 20 glucose, 10 glutamine, at pH 7.0). Brains were quickly transferred to 10 μL of the same saline in a 200 μL microcentrifuge tube kept at 2-3°C in a Boekel Scientific TropiCooler Benchtop Incubator (model 260014, Fisher Scientific, Ottawa, ON, Canada), and once 30 brains were collected (2-2.5 min per brain, ∼ 70 min total), samples were flash frozen in liquid N_2_ and stored at −80°C until use.

### Brain Na^+^/K^+^-ATPase activity

Samples were thawed on ice and mixed with 140 μL SEID buffer (in mM: 150 sucrose, 10 EDTA, 50 imidazole, and 0.1% w/v deoxycholic acid) to a volume of 150 μL. Next, samples were transferred to a 1.7 mL microcentrifuge tube by centrifuging them upside-down inside the 1.7 mL tube at 2000 × *g* for 1 min and mixed with a further 100 μL of SEID buffer to a final volume of 250 μL (if any brains couldn’t be transferred, this was noted). Here, samples were first ground with a polypropylene pestle and secondly sonicated (Model Q55, QSonica Sonnicators, Newtown, CT, USA) twice for 5-6 s with a 25-30 s break on ice to avoid overheating. Lastly, crude homogenates were vortexed, split into five aliquots, and stored at −80°C until use.

To normalize Na^+^/K^+^-ATPase activity to protein content, we first ran a Bradford assay on one of the five aliquots: In a clear 96-well plate, 5 μL of crude homogenate and protein standards (0 to 2 mg mL^-1^ bovine serum albumin, ALB001.25, Bioshop Canada, Burlington, ON, CA) was loaded into wells in triplicate followed by 250 μL of Bradford reagent (B6916, Sigma Aldrich, St. Louis, MI, USA). After incubating for 10 min, the absorbance of both samples and standards were measured in a microplate spectrophotometer (BioTek Epoch, Vermont, USA) at 595 nm, and sample protein contents were calculated by reference to the standards.

Activity of the Na^+^/K^+^-ATPase was measured in the remaining four aliquots using a continuous, spectrophotometric assay at 5, 10, 15, and 25°C following the approach of McCormick and Bern (1989), modified by Jonusaite et al. (2011), and following the best approaches for optimizing activity (Moyes et al., 2021): Solution A (4 units mL^-1^ lactate dehydrogenase (L2500, Sigma), 5 units mL^-1^ pyruvate kinase (P1506, Sigma), 2.8 mM phosphoenolpyruvate (cyclohexylammonium salt, P3637, Sigma), 3.5 mM ATP (AD0020, Bio Basic, Markham, ON, Canada), 0.3 mM NADH (NBO642, Bio Basic), and 50 mM imidazole, pH = 7.5) was prepared and activated by mixing with a salt solution (in mM: 189 NaCl, 42 KCl, 10.5 MgCl_2_, and 50 mM imidazole) at a ratio of 3:1 (A to salt) to final salt concentrations of 47.25 mM NaCl, 10.5 mM KCl, 2.63 mM MgCl_2_. Next, the quality of solution A was tested by creating a standard curve to describe the disappearance of NADH given the addition of specific amounts of ADP (to simulate hydrolysis of ATP to ADP by the Na^+^/K^+^-ATPase). This was done by adding known amounts of ADP (AD0016D, Bio Basic) in 10 μL aliquots (in triplicate) to a clear 96-well plate, followed by 200 μL of solution A, after which the depletion of NADH was measured at 340 nm every minute for 30 min in the microplate spectrophotometer. If a negative relationship between final absorbance (ABS) and ADP addition equal to or steeper than −0.013 ABS nmol ADP^-1^ was observed, solution A was deemed satisfactory (see an example in **Fig. S1**), and an accompanying solution B was created as described above with the addition of 5 mM ouabain (to specifically inhibit the Na^+^/K^+^-ATPase). To measure sample Na^+^/K^+^-ATPase activity, four 10 μL aliquots of sample homogenate were loaded into a clear 96-well plate followed by the addition of 200 μL of solution A to two wells with 200 μL of solution B to the remaining two, and again the depletion of NADH was monitored in the microplate spectrophotometer at 340 nm for 30 min. Activity of the Na^+^/K^+^-ATPase was calculated as follows:

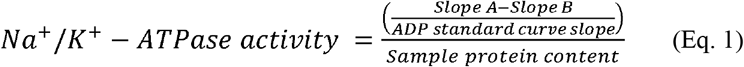

*Slope A* and *Slope B* refers the slope of disappearance of NADH (i.e. production of ADP) in the sample wells (*A*) and sample wells with the Na^+^/K^+^-ATPase blocked (*B*) and the *ADP standard curve slope* being the slope of the relationship between ADP addition and NADH disappearance (−0.013 and −0.015 ABS nmol ADP^-1^). Slopes of NADH disappearance in samples (*A* and *B*) were calculated on a linear part of the reading lasting at least 10 minutes. The initial part of readings was often not linear when temperature of the plate was still stabilizing (∼ 1-3 min). Eight biological replicates were measured for both warm- and cold-acclimated flies.

Thermal sensitivity of the Na^+^/K^+^-ATPase was estimated by calculating the Q_10_ values using the Q10() function from the ‘*respirometry*’ package in R (Birk, 2018), which fits an exponential curve and subsequently calculates Q_10_ using the formula:

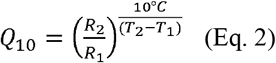

where R_2_ and R_1_ denote Na^+^/K^+^-ATPase activities at temperatures T_2_ and T_1_, respectively, with T_2_ being the higher temperature. This calculation was applied to individual samples (each sample was measured at all four temperatures), which allows us to statistically compare the thermal sensitivities of Na^+^/K^+^-ATPases in the brains of 15°C and 25°C acclimated flies.

### Brain Na^+^/K^+^-ATPase abundance

Abundance of the Na^+^/K^+^-ATPase was compared between warm- and cold-acclimated flies using Western blots. Samples were dissected and stored as described for the Na^+^/K^+^-ATPase activity assay above but with only ten brains per sample. Samples (10 μL) were thawed on ice and mixed with 90 μL RIPA buffer (50 mM Tris-HCl, 150 mM NaCl, 0.1% SDS, 1% deoxycholic acid, 1% Triton-X-100, 1 mM DTT, 1 mM PMSF, and 1:200 protease inhibitor cocktail (P1860, Sigma-Aldrich), pH 7.5) and transferred to a 1.7 mL microcentrifuge tube and homogenized as described above. Sample homogenates were centrifuged at 10000 × g for 10 min and supernatant was transferred to 200 μL tubes and kept on ice before sample protein contents were estimated using a Bradford assay (also described above). Before running samples through SDS-PAGE, a pilot experiment was run (using the procedure described below) to determine the amount of sample protein to load to optimize signal-to-noise ratios and avoid oversaturation of the final blot (Bass et al., 2017). Ultimately, protein samples were diluted with incomplete RIPA buffer (RIPA without DTT, PMSF, and protease inhibitor) and added to a 6 × Laemmli loading buffer (375 mM Tris-HCl, 9% SDS, 50% glycerol, and 0.03% bromophenol blue mixed 9:1 with β-mercaptoethanol) such that a total of 2 μg protein in 20 μL of buffer was loaded into wells of a Mini-PROTEAN TGX Stain-Free gel (5-14%) (Bio-Rad, Mississauga, ON, Canada). Protein ladder (5 μL of Precision Plus Protein™ All Blue Prestained Protein Standards, BioRad) was added to wells at the left and right of protein samples. Next, proteins were separated by running the gel in an electrophoresis chamber (Mini-PROTEAN Tetra Vertical Electrophoresis Cell, Bio-Rad) at 200 V for 30 min. Following electrophoresis, the gel was extracted and imaged on a ChemiDoc MP Imaging System (Bio-Rad) to confirm protein separation, after which the separated protein was transferred to a polyvinylidene fluoride (PVDF) membrane in a Trans-Blot Turbo System (Bio-Rad) at 25 V (1.3 A) for 7 min. Before blocking the PVDF membrane, it was imaged in the ChemiDoc MP Imaging System as the use of TGX Stain-Free gel allowed for imaging of membrane-bound total protein without subsequent staining (setting: Stain-free blot, 7.5 s exposure time). After imaging, the PVDF membrane was blocked in a 5% skim milk solution (w/w in TBS-T buffer; 20 mM Tris, 150 mM NaCl, 0.1% Tween-20 (v/w)) at room temperature (22-23°C) for 1 h. Blocked membranes were briefly rinsed in TBS-T buffer (3 × 5 min) and probed in a 500:1 solution of blocking buffer and primary antibody targeting the α-subunit of the Na^+^/K^+^-ATPase (mouse anti-Na^+^/K^+^-ATPase; a5, Developmental Studies Hybridoma Bank, Iowa City, IA, USA) at 4°C for 20 h overnight on a rotating stage. The next morning, probed membranes were washed in TBS-T (3 × 5 min) and incubated with the secondary antibody (StarBright Blue 700 Goat Anti-Mouse IgG) in blocking buffer (1:3000) at room temperature for 1 h on a rotating stage and subsequently washed (3 × 15 min) in TBS-T. Lastly, the blotted membrane was imaged in the ChemiDoc MP Imaging System to assess the fluorescent signal of the secondary antibody (setting: Starbright B700, 3 s exposure time). Densitometric quantification of total protein load (for normalization) and fluorescent signal was performed by measuring lane and band intensities, respectively, using Image Lab software (v6.0.1, Bio-Rad) (analysis files are attached as **Supplementary data**). Six biological replicates were measured for each acclimation group.

### Statistical analyses

All statistical analyses were performed in R version 4.0.2 (R Core Team, 2019). All data sets were tested for normality with a combination of Shapiro-Wilk tests and inspection of boxplots and residuals. Where the normality was violated, non-parametric tests were used to investigate differences between acclimation groups.

SD temperature and amplitude were confirmed to have similar variances using F tests, after which the means were compared using Student’s t tests. This was followed by a linear regression to investigate the relationship between the temperature and amplitude of the SD events. Descending slopes of the SD events we found to have different variances between the two acclimation groups, and means were therefore compared using a Welch t test. Subsequently, a linear regression was used to investigate the relationship between temperature and slope of the events.

To describe the relationship between temperature and reliance on the Na^+^/K^+^-ATPase (i.e. Latency) we used the nls() function to fit a three-parameter Gaussian model to the dataset on the effect of temperature on the latency to ouabain-induced SD using the following formula:

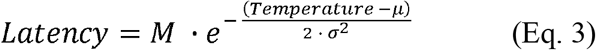

where *M* is the maximum latency (in min), *μ* is the temperature where latency is maximized (in °C), and *σ* is the ‘standard deviation’ (in °C) and can be used to represent a measure of ‘thermal insensitivity’ such that a higher value denotes a lower effect of temperature (i.e. in this Gaussian fit, *σ* represents the temperature change needed to cause a ∼ 39.3 % drop from the maximum, rather than representing an area under the curve in the traditional sense of Gaussian ‘standard deviation’). To compare the effect of acclimation temperature on latency to SD between cold- and warm-acclimated flies, we expanded the outputs of the fitted Gaussian models (*M, μ, σ*) to include acclimation terms (*a*_*i*_, where *i* denotes the parameter in question) for each of the three above mentioned parameters (i.e. [*M* + *a*_*M*_ ·Acclimation temperature], [*μ* + *a*_*μ*_ · Acclimation temperature], and [*σ* + *a*_*σ*_ · Acclimation temperature]):

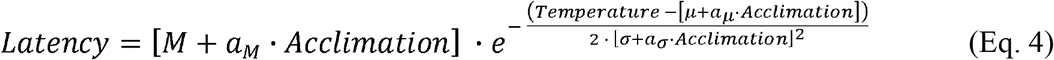

This procedure allows us to use the nls() function to determine if and how *M, μ*, and *σ* were each affected by thermal acclimation using non-linear least squares regression. A summary of the output parameters (*M, μ, σ, a*_*M*_, *a*_*μ*_, *a*_*σ*_) and statistics (t- and P-values) can be found in **Table 1**.

**Table 1.** Non-linear least squares regression to a Gaussian model was used to estimate the effect of acclimation on the thermal performance curve of the latency to spreading depolarization dataset. The output parameters from the model include the maximum latency (*M*), the temperature of the maximum latency (*μ*, midpoint of distribution), a measure of the thermal sensitivity (*σ*, the standard deviation), and accompanying acclimation parameters (*a*_*M*_, *a*_*μ*_, and *a*_*σ*_) to estimate how acclimation affected the main parameters (see the *Statistical analyses* section for more details).

| Main parameter | Model output | Acclimation parameter | Effect of acclimation | t | P | Acclimation group estimates |  |
| --- | --- | --- | --- | --- | --- | --- | --- |
|  |  |  |  |  |  | 15°C | 25°C |
| $M$ | $56.0 \pm 12.3$ min | $a_M$ | $1.0 \pm 0.6$ min °C<sup>-1</sup> | 1.6 | 0.115 | $70.6 \pm 4.3$ min | $80.4 \pm 4.3$ min |
| $\mu$ | $17.1 \pm 1.7$ °C | $a_\mu$ | $0.4 \pm 0.1$ °C °C<sup>-1</sup> | 5.9 | < 0.001 | $23.6 \pm 0.6$ °C | $28.0 \pm 0.4$ °C |
| $\sigma$ | $10.2 \pm 1.9$ °C | $a_\sigma$ | $-0.2 \pm 0.1$ °C °C<sup>-1</sup> | -2.0 | 0.050 | $7.7 \pm 0.7$ °C | $6.0 \pm 0.4$ °C |

The total sample size of 68 results in 62 degrees of freedom for the analysis.

The effect of temperature on the activity of brain Na^+^/K^+^-ATPases in cold- and warm-acclimated flies was analyzed using a linear mixed-effect model, where assay temperature and acclimation temperature were treated as fixed factors, while samples were included as random factors (because the same biological replicates were tested at all temperatures). Here, Q_10_ was calculated for each sample to estimate the thermal sensitivity of the brain Na^+^/K^+^-ATPase in both groups 1) across the entire temperature spectrum (5-25°C), and 2) in the three temperature intervals (5-10°C, 10-15°C, and 15-25°C) using the Q10() function from the ‘*respirometry*’ package (Birk, 2018). Q_10_ across the entire spectrum was compared between groups using a Mann-Whitney-Wilcoxon U test as the data were not normally distributed, while the Q_10_ values in all intervals was analyzed using a linear mixed effect model with temperature and acclimation as fixed factors and the individual sample as a random factor. Q_10_ values were found to vary greatly depending on the temperature interval investigated and Na^+^/K^+^-ATPase activity did not appear to display an exponential relation with temperature for either group. We therefore decided to further analyze this dataset similar to how we investigated the effect of acclimation on the latency to ouabain-induced spreading depolarization. Thus, we fitted the three-parameter Gaussian model (Eq. 3) to Na^+^/K^+^-ATPase activity for each sample which allowed us to more accurately investigate and compare maximum activity (*M*), temperature of maximum activity (*μ*), and thermal sensitivity (*σ*) of the two groups. Here, group parameters were compared using a Mann-Whitney-Wilcoxon U test for *M* (one group was non-normal) and Welch t tests for *μ* and *σ* (F tests revealed that group variation estimates differed).

Lastly, we compared the abundance of Na^+^/K^+^-ATPase in the brains of cold- and warm-acclimated flies using Western blots. The fluorescent signal was normalized to total protein load for each sample, and all samples were subsequently normalized to the mean of the 25°C acclimated group. Data were log-transformed during analysis, and groups were compared using a linear mixed effect model with acclimation temperature as a fixed factor and blot as a random factor (as samples were measured on different blots). A significance level of 0.05 was applied to all analyses.

## Results

### Cold acclimation lowers spreading depolarization temperature, amplitude, and descend slope

As expected, cold acclimation reduced SD temperature to 5.1 ± 0.4°C compared to 9.0 ± 0.4°C after warm acclimation (t_18_ = −7.1, P < 0.001, **Fig. 1A**). Similarly, cold acclimation led to a reduced amplitude of the SD event (t_18_ = −3.9, P = 0.001) such that the event had an amplitude of 29.9 ± 1.5 mV and 39.3 ± 1.9 mV in cold- and warm acclimated flies, respectively (**Fig. 1B**). Interestingly, there was a strong linear relationship between SD temperature and amplitude within individuals (t_17_ = 2.9, P = 0.010), and the slope of this relationship was not affected by acclimation (t_17_ = −0.1, P = 0.911) (**Fig. 1C**).

**Figure 1.**
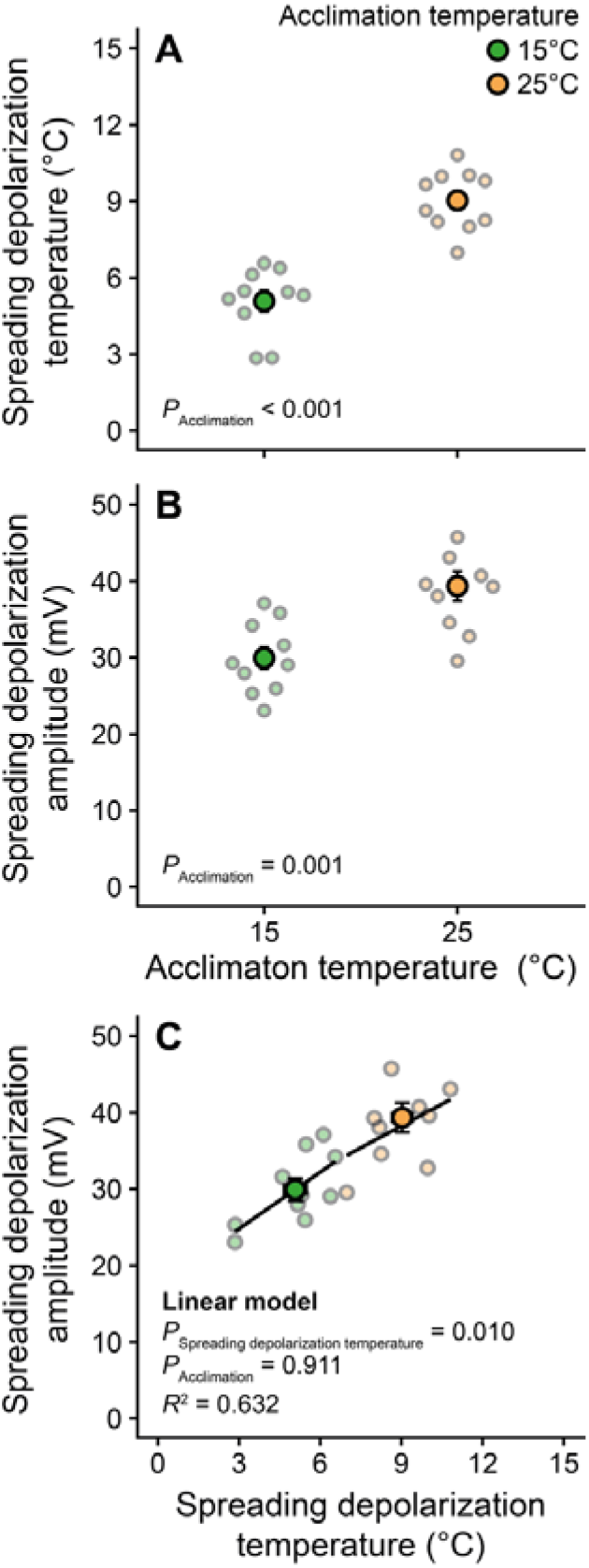
Cold acclimation lowers spreading depolarization temperature and amplitude. (A) Cold acclimation (15°C; green) of female *Drosophila melanogaster* lowers the spreading depolarization temperature relative to warm acclimation (25°C; orange). (B) The amplitude of the spreading depolarization event is similarly lowered by cold acclimation. (C) A statistically significant and positive relationship between spreading depolarization temperature and amplitude was found among individual flies, but there was no effect of acclimation temperature on the slope of the relationship. N = 10 flies per group, which are depicted by smaller, translucent points. Error bars not shown are obscured by the symbols.

The last SD parameter we characterized was the slope of the descending down-shift in transperineurial potential (**Fig. S2**). Here we found a clear trend that cold-acclimation had resulted in more gentle descend slopes (**Fig. S2A**), but while this trend was almost two-fold it did not reach the level of statistical significance (−6.4 ± 1.1 mV s^-1^ and −11.3 ± 2.4 mV s^-1^ for cold- and warm-acclimated flies, respectively; t_12.8_ = 1.9, P = 0.085). Furthermore, we noted that SD events occurring at a lower temperature tended to have gentler slopes in general (**Fig. S2B**), but this too failed to reach statistical significance (t_18_ = −2.0, P = 0.062).

### Cold acclimation left-shifts the thermal sensitivity of ouabain-induced spreading depolarization

To investigate how acclimation affected the role of Na^+^/K^+^-ATPase in preventing spreading depolarization, we injected the specific Na^+^/K^+^-ATPase inhibitor ouabain into the heads of flies while monitoring the transperineurial potential. Ouabain injection always induced one or more spreading depolarization events (see **Fig. 2** for example trace), and we used the latency to the first event as a measure of how the event relates to Na^+^/K^+^-ATPase. The overall pattern resembled a thermal performance curve (**Fig. 3**), so we decided to use a model-fitting approach to analyze these data (see *Statistical analyses*). Fitting a three-parameter Gaussian model to data from the two groups resulted in high correlation coefficients (R^2^) of 0.686 and 0.801 for cold- and warm acclimated flies, respectively (**Eq. 3**), and 0.747 for the combined model (**Eq. 4**). Using the Gaussian model outputs to statistically compare the effect of cold- and warm-acclimation revealed that while cold-acclimated flies had a similar maximum latency (*M*) compared to their warm-acclimated conspecifics (*a*_*M*_; t_62_ = 1.6, P = 0.115), the midpoint of the curve (μ) of cold-acclimated flies was left-shifted by 4.3 ± 0.7 °C (*a*_*μ*_; t_62_ = 5.9, P < 0.001) and a thermal ‘insensitivity’ factor (σ) was increased by 1.7 ± 0.8°C (*a*_*σ*_; t_62_ = −2.0, P = 0.050; **Fig. 3**). All model parameters and statistical outputs from this analysis can be found in **Table 1**.

**Figure 2.**
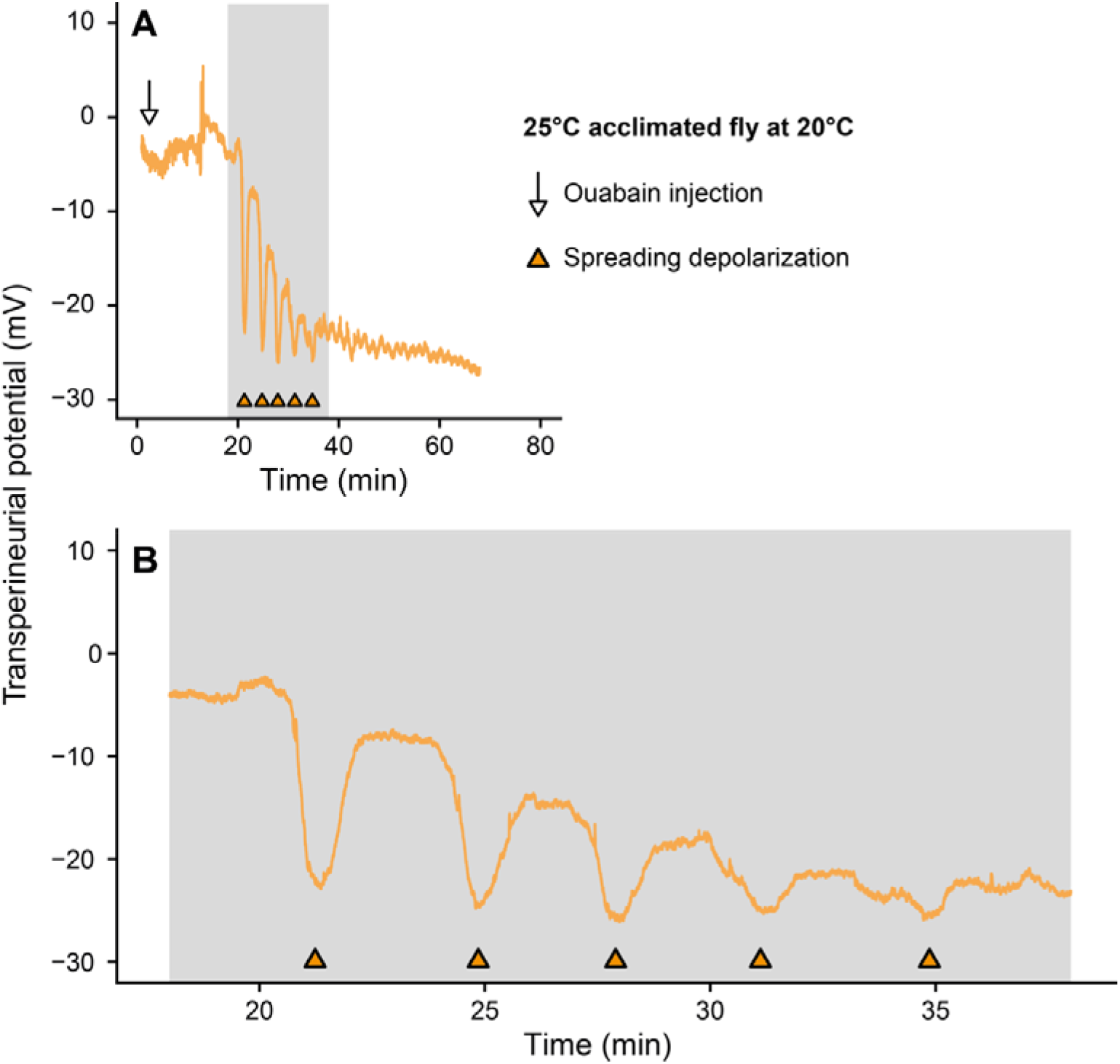
Example trace of spreading depolarization events caused by ouabain injection into the head of *Drosophila melanogaster*. (A) Injection of the Na^+^/K^+^-ATPase-specific blocker ouabain (arrow) leads to spreading depolarization in the brain (triangles) after a delay. (B) A closer look at the shaded area in panel A, showing the spreading depolarization events, which are characterized by large, rapid negative shifts in the transperineurial potential. In the example depicted, ouabain injection elicited repeated (five) spreading depolarization events, but generally the number of events was positively related to temperature (see **Fig. S3**). Note that the triangles here point to the transperineurial potential minima of the SD events and not their half-amplitude which was used in latency estimates.

**Figure 3.**
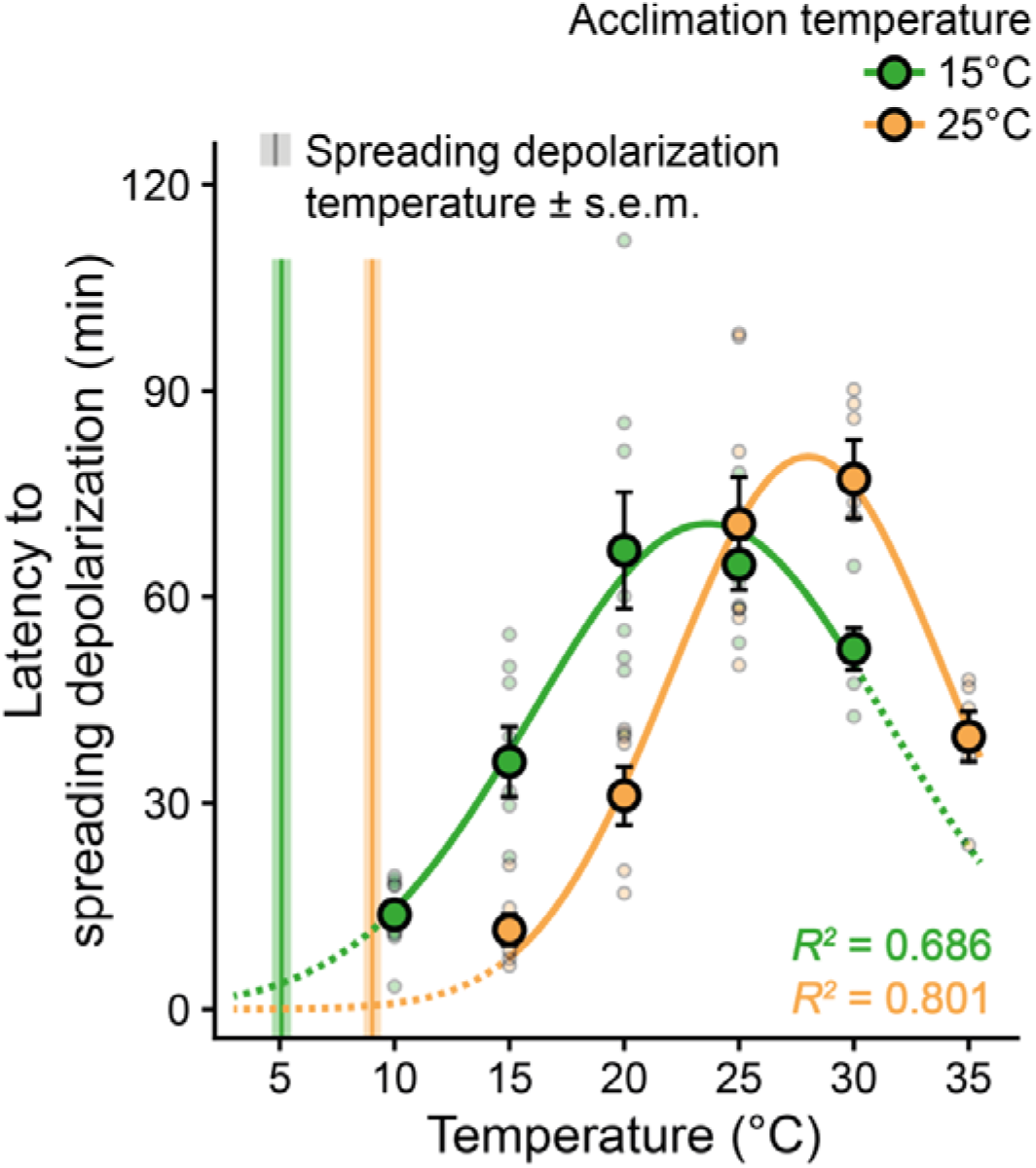
Cold acclimation causes a leftward thermal-shift in the latency to ouabain-induced spreading depolarization in the brain. Latency to spreading depolarization after ouabain injection was highly temperature-dependent, and this effect of temperature was left-shifted in cold-acclimated (15°C; green) relative to warm-acclimated (25°C; orange) flies. The thermal performance curve analysis is left-shifted in the cold-acclimated group (parameter analysis in **Table 1**) and latency approached zero (extrapolation) around the temperature found to cause spreading depolarization during gradual cooling (*sensu* **Fig. 2A**; vertical lines and shaded areas depict means and standard errors). N = 6-8 per temperature and acclimation group. Some data points and error bars are obscured by the symbols. Dotted lines represent model extrapolations beyond the data.

### Thermal kinetics of brain Na^+^/K^+^-ATPase are altered by cold acclimation

Activity of the Na^+^/K^+^-ATPase was measured in at 5, 10, 15, and 25°C in brain samples for each cold- or warm-acclimated brain sample (**Fig. 4**). Overall, there was no main effect of acclimation on the activity of Na^+^/K^+^-ATPase in the brain (F_1,42_ = 1.5, P = 0.248). Activity of the pump was highest at 25°C in both acclimation groups with warm-acclimated flies having higher activity (38.0 ± 0.9 μmol ADP mg^-1^ h^-1^) than their cold-acclimated conspecifics (31.4 ± 1.1 μmol ADP mg^-1^ h^-1^) (**Fig. 4A**). As temperature was lowered, the activity of the Na^+^/K^+^-ATPase decreased (F_3,42_ = 1039, P < 0.001), but in an acclimation-specific manner. The magnitude of the temperature effect was significantly mitigated by cold acclimation (F_3,42_ = 29.7, P < 0.001), resulting in cold-acclimated flies having higher brain Na^+^/K^+^-ATPase activity at 5°C (6.9 ± 0.5 μmol ADP mg^-1^ h^-1^) than the warm-acclimated flies (4.8 ± 0.2 μmol ADP mg^-1^ h^-1^) (**Fig. 4A**). The smaller effect of cold on brain Na^+^/K^+^-ATPase activity in cold-acclimated flies resulted in a Q_10_ of 1.83 ± 0.05 across the entire temperature range which was significantly smaller than the 2.22 ± 0.04 observed in warm-acclimated flies (Mann–Whitney-Wilcoxon *U* = 1, N = 16, P < 0.001) (**Fig. 4B**). Calculating Q_10_ in each temperature interval and for every sample revealed a similarly strong effect of acclimation on the temperature sensitivity of the brain Na^+^/K^+^-ATPase (F_1,28_ = 39.1, P < 0.001), however, this analysis also revealed that Q_10_ for both acclimation groups increased at lower temperatures (F_2,28_ = 105, P < 0.001) and that the degree to which it increased was smaller after cold acclimation (F_1,28_ = 7.2, P = 0.003). Thus, the relation we observed between temperature and brain Na^+^/K^+^-ATPase activity appeared non-exponential within the temperature range of this study, and we therefore applied a Gaussian fit to each sample to more accurately describe the relation and compare acclimation groups (see **Fig. S3**). Using this approach we found no evidence of a shift in the Na^+^/K^+^-ATPase activity performance curve (*μ*; t_7.3_ = 1.2, P = 0.251), but instead that the cold-acclimated flies had a lower maximum brain Na^+^/K^+^-ATPase activity (*M*; Mann–Whitney-Wilcoxon *U* = 12, N = 16, P = 0.038) and a higher temperature ‘insensitivity’ (*σ*; t_7.1_ = 2.4, P = 0.047) (**Fig. S3**). Thus, both analysis approaches suggest that cold acclimation lowers the thermal sensitivity of Na^+^/K^+^-ATPase.

**Figure 4.**
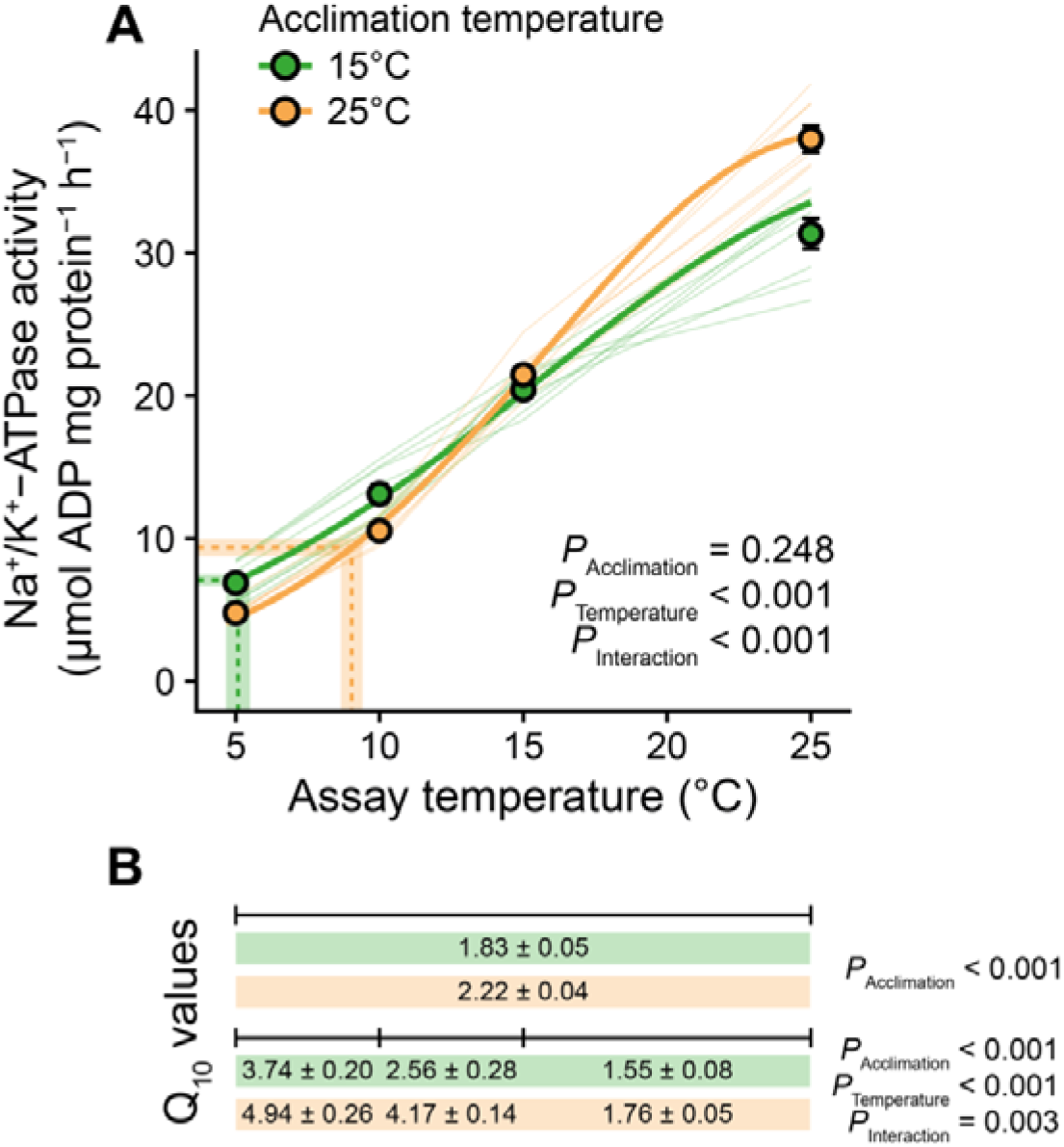
Cold acclimation lowers the *in vitro* thermal sensitivity of Na^+^/K^+^-ATPase extracted from the brain of *Drosophila melanogaster*. (A) The activity of Na^+^/K^+^-ATPase from brains of cold-acclimated flies (green) did not differ in their overall activity when compared to warm-acclimated flies (green), but their Na^+^/K^+^-ATPases were less sensitive to temperature such that their activity was higher at 5°C. Thin lines depict model outputs for each biological replicate (N = 8 per acclimation group), and solid circles denote group means at each temperature. *In vitro* pump activity at the SD temperature (based on calculations using Gaussian model parameters; dashed lines and shaded areas denote group means and errors, respectively) was slightly lower in the cold-acclimated group. (B) The difference in pump thermal sensitivity between the two acclimation groups is evident from Q_10_ values calculated for each sample across the measurement temperature range (see Methods for details). The Q_10_ of Na^+^/K^+^-ATPase activity of cold-acclimated flies is consistently lower than warm-acclimated flies, regardless of the temperature interval tested. The Q_10_ estimates are, however, not consistent across the range of temperatures measured here, and we therefore used a modelling approach similar to the one used in **Table 1** which also confirmed the lowered thermal sensitivity (see **Fig. S3**).

### Acclimation does not change the abundance of brain Na^+^/K^+^-ATPase

To investigate if cold- and warm-acclimated flies differed in Na^+^/K-ATPase content, we measured Na^+^/K-ATPase abundance using Western blots targeting the α subunit (example blot in **Fig. 5A and 5B**). We found no difference in brain Na^+^/K^+^-ATPase abundance between cold- and warm-acclimated flies (t_8_ = 0.1, P = 0.922) (**Fig. 5C**).

**Figure 5.**
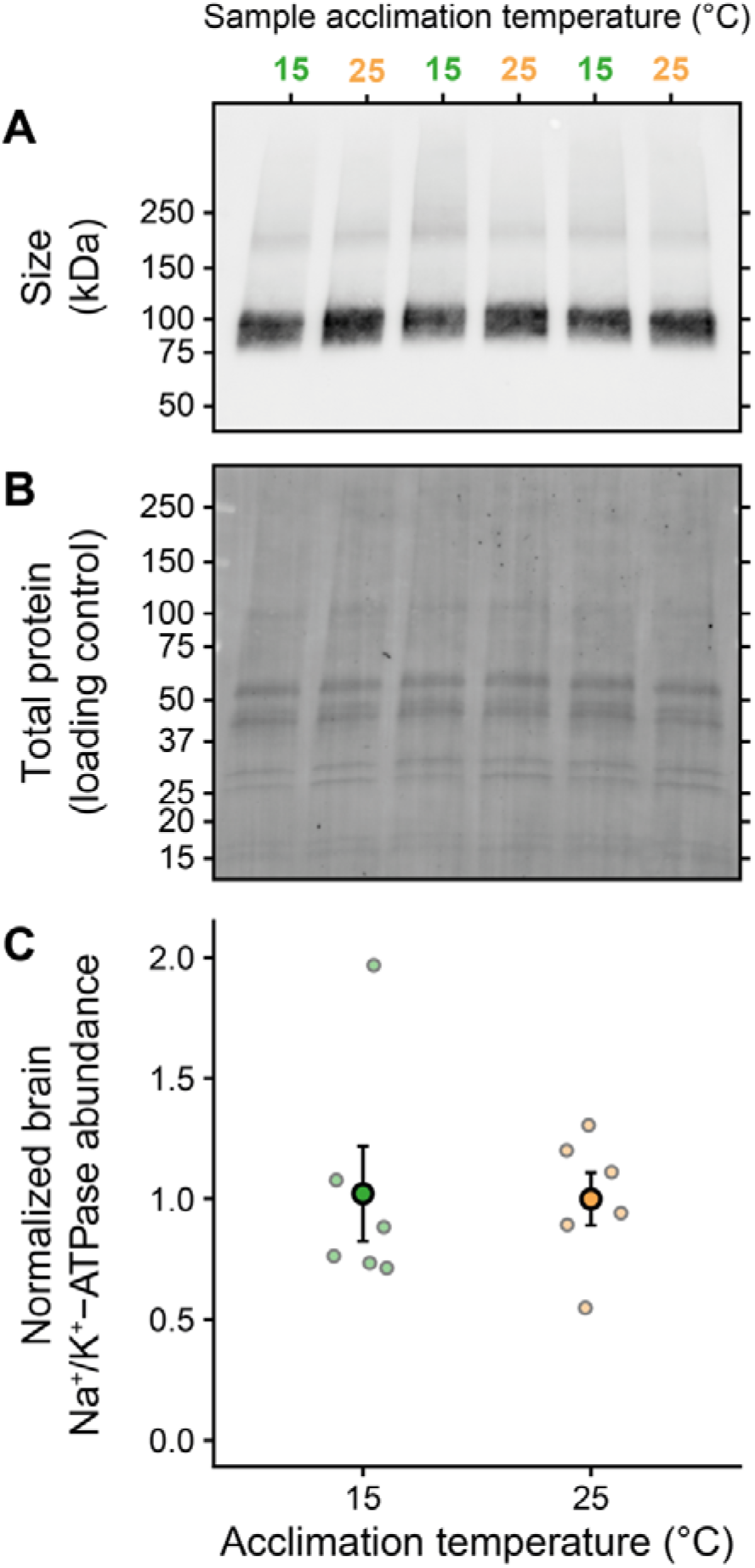
Acclimation does not alter the abundance of the α subunit of Na^+^/K^+^-ATPase in the brain of *Drosophila melanogaster*. (A) Example blot showing a prominent band at the expected size of ∼ 105 kDa. (B) Matching stain free total protein blot used for normalizing the fluorescent signals from the bands in panel A to total protein load. (C) Comparing the Na^+^/K^+^-ATPase α-subunit abundance (normalized to the 25°C acclimated group) revealed no difference between the groups. N = 6 per group.

## Discussion

Thermal tolerance is broadly considered to be important in determining the distribution of insects (Addo-Bediako et al., 2000; Kimura, 2004; Sunday et al., 2011; Kellermann et al., 2012), and the capacity to respond and adapt to environmental change is therefore likely to shape their future biogeography (Chown and Nicolson, 2004; Ghalambor et al., 2007; Overgaard et al., 2011). In many insects, including *Drosophila*, cold acclimation has repeatedly been shown to induce adaptive improvements in the ability to maintain mobility at low temperatures (Kelty and Lee Jr, 1999; Ransberry et al., 2011; Schou et al., 2017). Here, we reaffirm this cold acclimation capacity and argue that modifications to a single ion transporter, the Na^+^/K^+^-ATPase, is critical to cold acclimation in the brain of *Drosophila* and reduces the CT_min_ by lowering the temperature at which a spreading depolarization shuts down central nervous system function.

### Maintenance of Na^+^/K^+^-ATPase activity decreases spreading depolarization susceptibility

A central theme in the current model of insect cold tolerance is that more cold tolerant insects better maintain function of active ion transporters at low temperature, which allows them to maintain ion balance and avoid tissue damage induced by hyperkalemia (Overgaard and MacMillan, 2017; Overgaard et al., 2021). In particular, the Na^+^/K^+^-ATPase is central to maintenance of ion and water balance in the insect central nervous system (Treherne and Schofield, 1981). We therefore chose to investigate the role of the Na^+^/K^+^-ATPase in cold-induced SD by 1) inhibiting the Na^+^/K^+^-ATPase with ouabain to estimate sensitivity to SD *in vivo* and 2) measuring the activity of the Na^+^/K^+^-ATPase *in vitro* at benign and low temperatures after both warm- and cold acclimation.

Application of ouabain to the brain *in vivo* resulted in SD after a delay, and this latency was both acclimation- and temperature-specific, following the general shape of a thermal performance curve that was left-shifted and had a lower thermal sensitivity in the cold-acclimated flies (**Fig. 3**; **Table 1**). The magnitude of the shift in latency (4.3 ± 0.7 °C) closely matches that of the shift in SD temperature (4.0 ± 0.6 °C), indicating a strong link between SD and Na^+^/K^+^-ATPase activity.

Ouabain specifically blocks the Na^+^/K^+^-ATPase and the latency to SD after ouabain-injection could therefore be interpreted as a measure of ‘reliance’ of the brain on the Na^+^/K^+^-ATPase in maintaining function (i.e. prevent SD). We found that cold-acclimated flies are able to ‘rely’ on the activity of the Na^+^/K^+^-ATPase down to a lower temperature, likely resulting in a lower organismal CT_min_. One consideration is the manner in which ouabain binds to the Na^+^/K^+^-ATPase. Specifically, it has been demonstrated that the binding of ouabain to the Na^+^/K^+^-ATPase is temperature-sensitive indicating that the Na^+^/K^+^-ATPase needs to be active to bind ouabain (Clausen and Hansen, 1974; Bayley et al., 2020). This means that the thermal performance curve we observe here could be a product of the activity of the Na^+^/K^+^-ATPase itself (see **Fig. 3**). However, when looking at the direct effect of temperature on Na^+^/K^+^-ATPase activity measured *in vitro* the pattern is different; while the thermal sensitivity is still lower in the cold-acclimated group, there is no evidence for a left-shift in the performance curve, and the maximum activity is lower in the cold-acclimated group (**Fig. 4** and **Fig. S3**), indicating that the differences between acclimation groups may not be directly related to the innate activity of the Na^+^/K^+^-ATPase, but rather to its’ activity in the native membrane environment and/or cellular location. The emerging thermal performance curves could also be a product of the thermal sensitivity of compensatory ionoregulatory mechanisms independent of the inhibited Na^+^/K^+^-ATPase. Such mechanisms would need to be directly affected by temperature to produce a thermal performance curve, making other active transporters likely candidates (for example the V-type H^+^-ATPase, which can modulate the shape of the SD event (see Robertson and Van Dusen (2021)). Regardless of the underlying mechanism, our results support previous observations of an increased latency to ouabain-induced SD in more cold tolerant insects (Gantz et al., 2020). Similarly, application of ouabain has been shown to increase the susceptibility to SD during heating (Rodgers et al., 2007), and insects that are expected to be less temperature tolerant, be it by old age or experimental manipulation of ionoregulatory capacity, show the same trend (Spong and Robertson, 2013; Spong et al., 2014; Spong et al., 2015; Spong et al., 2016b).

Measuring the *in vitro* activity of the Na^+^/K^+^-ATPase in homogenized samples revealed a strong effect of temperature, which was affected by acclimation such that the negative effect of cold was smaller after cold acclimation (**Fig. 4**). These findings largely reflect the observations of Cheslock et al. (2021) who measured Na^+^/K^+^-ATPase in whole-head samples. The temperature-dependency of Na^+^/K^+^-ATPase activity and the effect of acclimation were also evident from the Q_10_ values, which were lower in the cold-acclimated group of flies (**Fig. 4B**), despite our Q_10_’s being low compared to previous estimates for invertebrate Na^+^/K^+^-ATPases (Leong and Manahan, 1997; Pan et al., 2016). However, our data suggest that the relationship between Na^+^/K^+^-ATPase activity and temperature within our range of temperatures is non-exponential and that applying a single Q_10_ across all temperatures is inappropriate. Indeed, when we calculate Q_10_ at each temperature interval we find that Q_10_ itself is temperature-dependent (**Fig. 4B**). We therefore decided to try the three-parameter Gaussian function to model thermal performance curve parameters for *in vitro* activity as well. Similar to the Q_10_-approach the Gaussian model revealed a lowered thermal sensitivity for cold-acclimated flies (**Fig. S3**), but instead of showing a shift in optimum temperature, like the *in vivo* experiment on latency, it showed a lowered maximum activity at the optimal temperature (**Fig. S3**). The Gaussian model was, however, also suboptimal at describing the relationship between temperature and activity in our dataset as it partially relied on extrapolation beyond our data. Regardless, no matter the approach our data suggests that acclimation changes the thermal kinetics of the Na^+^/K^+^-ATPase. Numerous studies from the 1960’s and 1970’s investigated how enzymes were adapted to their thermal environment (see reviews by Hazel and Prosser (1974) and Hochachka and Somero (1973)), but a role of electrogenic ion transporters in the chill coma phenotype was only suggested later (see Goller and Esch (1990) and Hosler et al. (2000)), while the link between insect coma and SD was established even more recently (Robertson et al., 2017; Andersen et al., 2018). We demonstrate here how differences in the SD temperature may specifically relate to variation in the thermal dependency of the Na^+^/K^+^-ATPase. These findings are mainly correlative in nature, but they strongly suggest that the lowered chill coma temperature in cold-acclimated *D. melanogaster* is achieved *via* lowered thermal sensitivity of the active transporters involved in maintaining brain ion homeostasis (MacMillan and Sinclair, 2011). However, the specific mechanisms by which the brain Na^+^/K^+^-ATPase of cold-acclimated flies better maintains activity in the cold remain unknown.

Molecular mechanisms known to affect the activity of the Na^+^/K^+^-ATPase include posttranslational modifications, transcription of different Na^+^/K^+^-ATPase isoforms, changes in translation, and an altered membrane environment (Therien and Blostein, 2000): The insect Na^+^/K^+^-ATPase has several phosphorylation sites (Emery et al., 1998), and depending on the state of these sites the activity of the Na^+^/K^+^-ATPase can be greatly affected (Bertorello et al., 1991).

Phosphorylation state of the pump has been linked to seasonal improvements in cold tolerance, albeit in a freeze-tolerant species where it drives winter suppression of pump activity during dormancy (McMullen and Storey, 2008). *D. melanogaster* has one gene encoding the catalytic α-subunit and three genes encoding the regulatory β-subunit of the Na^+^/K^+^-ATPase (*ATPα* and *nrv1-3*, respectively) (Larkin et al., 2021). However, these genes all have several transcripts, which can likely be differentially translated, resulting in a suite of Na^+^/K^+^-ATPase isoforms that may vary in activity and function. Indeed, cold-adapted squid exhibit improved low temperature transport capacity and efficiency of a ubiquitously expressed Na^+^/K^+^-ATPase due to specific amino acid substitutions (Galarza-Muñoz et al., 2011). In mammals, there are multiple Na^+^/K^+^-ATPase isoforms that vary in activity and localization and reduced transport capacity in some isoforms, specifically when localized in astrocyte glia, leads to increased SD susceptibility (Clausen et al., 2016; Isaksen and Lykke-Hartmann, 2016; Clausen et al., 2017; Reiffurth et al., 2020). Thus, similar structural modifications to the Na^+^/K^+^-ATPase may underlie the variation in transport capacity and by extension susceptibility to SD (Andrew et al., 2022). Expression of specific Na^+^/K^+^-ATPase subunits has been investigated in relation to cold tolerance in *D. melanogaster* and a small increase in *nrv2* expression was noted after cold acclimation, albeit in whole animal samples (MacMillan et al., 2015b). Lastly, thermal acclimation or adaptation often results in altered physical properties of the cell membrane that act to defend fluidity during cold exposure (homeoviscous adaptation, see Hazel (1995) and Koštál (2010)), which is also true for *Drosophila* (Overgaard et al., 2005), and such changes in the membrane environment have been shown to impact transport capacity of the Na^+^/K^+^-ATPase (Else and Wu, 1999; Habeck et al., 2015). In our *in vitro* experiments the Na^+^/K^+^-ATPase was largely independent of the native membrane environment (Moyes et al., 2021), and our data therefore represent innate transport capacity, however, modifications to the membrane environment remain likely to affect Na^+^/K^+^-ATPase activity *in vivo*.

### Unaltered brain Na^+^/K^+^-ATPase abundance after acclimation indicates changes to localization

Abundance of the Na^+^/K^+^-ATPase α-subunit did not differ between cold- and warm-acclimated flies, confirming that the differences in activity measured in the Na^+^/K^+^-ATPase assay were not related to Na^+^/K^+^-ATPase content (**Fig. 5**). It therefore seems most likely that the difference in activity was driven by one of the many factors described above. This is the first time abundance of the Na^+^/K^+^-ATPase in the brain has been related to a specific measure of insect cold tolerance, but a study on whole-body Na^+^/K^+^-ATPase α-subunit abundance also reported no difference between cold- and warm-acclimated flies (MacMillan et al., 2015b), and a study on the physiology underlying improved heat tolerance following heat hardening in locusts also found no difference (Hou et al., 2014). Interestingly, Hou et al. (2014) found that localization of the Na^+^/K^+^-ATPase changed drastically during heat hardening as more Na^+^/K^+^-ATPases were incorporated into the membranes of neuronal somata, perineurial cells and nerve axons, and similar observation were made in relation to cold hardening more recently (Robertson and Moyes, 2022). In the present study we did not investigate localization, but we hypothesize that the reduced SD amplitude following acclimation could relate to increased Na^+^/K^+^-ATPase activity in the adglial membrane of perineurial and subperineurial glia, which would also serve to delay SD during cooling due to the maintained ionoregulatory capacity of the cold-acclimated Na^+^/K^+^-ATPases. Furthermore, we speculate that localization might be one of the key parameters responsible for the two different thermal performance curve shapes of the thermal dependency of the Na^+^/K^+^-ATPase (**Fig. 3 and 4**) as a key methodological difference between the *in vivo* and *in vitro* experiments is tissue homogenization. Thus, the thermal performance curve *in vivo* (**Fig. 3**) could represent reliance on (or activity of) the Na^+^/K^+^-ATPase when localized to native membranes, sensitivity of the Na^+^/K^+^-ATPase to ouabain, or a combination of thereof, which clearly suggests that 1) localization to the native cell membrane impacts activity, and 2) the left-shift caused by cold-acclimation is driven by a combination of temperature effects on the Na^+^/K^+^-ATPase and the membrane. This is, however, mainly speculation, and more research is needed to investigate the molecular and/or cellular mechanisms by which the Na^+^/K^+^-ATPase is able to maintain transport capacity in the cold after cold-acclimation.

### Thermal acclimation alters the nature of the spreading depolarization event

Developmental acclimation of flies to 15°C not only lowered the temperature threshold for cold-induced SD onset, it also decreased the amplitude and descend slope of the drop in transperineurial potential (see **Fig. 1** and **Fig. S2**). The transperineurial potential is the electrical potential generated across the insect blood brain barrier and is derived from the difference in electrical potential generated across the basolateral (facing the hemolymph) and the adglial (facing the neurons) membranes of the glia (transperineurial = basolateral - adglial) (Schofield and Treherne, 1984). The general consensus is that the large shift in transperineurial potential during SD reflects a near-complete depolarization of the adglial membrane (Robertson et al., 2020). The lower SD amplitude we observe in the cold-acclimated flies could therefore reflect 1) a less polarized adglial membrane before SD, 2) a more polarized membrane during SD, or 3) a combination of both. Similar to us, Cheslock et al. (2021) found a reduced SD amplitude after cold-acclimation in *D. melanogaster*, but they also found a more positive transperineurial potential at benign temperature, indicating a more polarized adglial membrane prior to SD. However, because both the basolateral and adglial membrane potentials affect the transperineurial potential it is difficult to determine the exact origin of the change caused by acclimation. Nonetheless, we speculate that ionoregulatory changes play a key role in determining the abovementioned differences in transperineurial potential and SD amplitude: First, Na^+^/K^+^-ATPase activity generates a considerable electrogenic potential in insects (Wareham et al., 1974; Bayley et al., 2021) and glial Na^+^/K^+^-ATPase localization is therefore highly likely to affect the transperineurial potential. Specifically, deployment of thermally insensitive Na^+^/K^+^-ATPases to the adglial membrane would tend to increase adglial polarization and drive the transperineurial potential up, as reported by Cheslock et al. (2021). Secondly, the smaller disruption of ion balance experienced by cold-acclimated flies during SD could leave the adglial membrane less depolarized (Andersen et al., 2018). Lastly, changes to the interstitial K^+^ clearance, glial K^+^ shuttling, and basolateral K^+^ efflux could be involved (i.e. glial spatial buffering (Orkand et al., 1966; Smith and Shipley, 1990; Kocmarek and O’Donnell, 2011; Chen and Swale, 2018). Indeed, the hypothesized deployment of Na^+^/K^+^-ATPases to the adglial membrane would facilitate glial K^+^ uptake, and could also partially explain the gentler descend slope in the transperineurial potential during the SD onset as an improved capacity to acutely buffer the initial increase in interstitial K^+^ would tend to slow the overall progress of the event. This is, however, mainly speculation, and more research is needed to investigate the potential links between SD amplitude, adglial and basolateral membrane potentials, Na^+^/K^+^-ATPase activity, and glial spatial buffering. Furthermore, the two acclimation groups experience SD at different temperatures and we can therefore not rule out direct effects of temperature on the membrane potentials (see **Fig. 1C and S2B**).

## Conclusion

In summary, we demonstrate that phenotypic plasticity in the CT_min_ of *D. melanogaster*, here estimated as the SD temperature, relates to the ability to maintain active transport capacity of the Na^+^/K^+^-ATPase in the brain at low temperature *in vivo* and *in vitro* without altering the overall abundance of the Na^+^/K^+^-ATPase. Considering the central role of the Na^+^/K^+^-ATPase in regulating extracellular ion homeostasis in the insect central nervous system it is perhaps not surprising that its activity is essential for preventing SD when exposed to stressful cold. Nonetheless, our findings support a key role of the Na^+^/K^+^-ATPase in modulating SD susceptibility, and adaptations promoting maintained active ion transport capacity therefore appear to be a key pathway by which cold tolerance can be improved at multiple levels of biological organization (see Overgaard et al. (2021)). The molecular and/or cellular mechanism by which the Na^+^/K^+^-ATPase of cold-acclimated flies is able to maintain activity at low temperature remains unknown, but we suggest that future research should focus on investigating the roles of phosphorylation, transcript expression, sequence variation, cellular localization, and the membrane environment. Lastly, we speculate that the observed changes to SD parameters (amplitude and slope) relate to improved ionoregulatory capacity at low temperature and a hypothesized deployment of Na^+^/K^+^-ATPases to the adglial membranes of the blood-brain barrier.

## Acknowledgements

None

## Author contributions

The study was conceived and designed by MKA and HAM. MKA performed the experiments, analyzed the data, and wrote the first draft. All authors made edits to the manuscript and approved the final version of the manuscript.

## Data availability

All data has been made available to reviewers during the review process and will be uploaded as supplementary material upon acceptance and publication.

## Supplementary material

**Figure S1.**
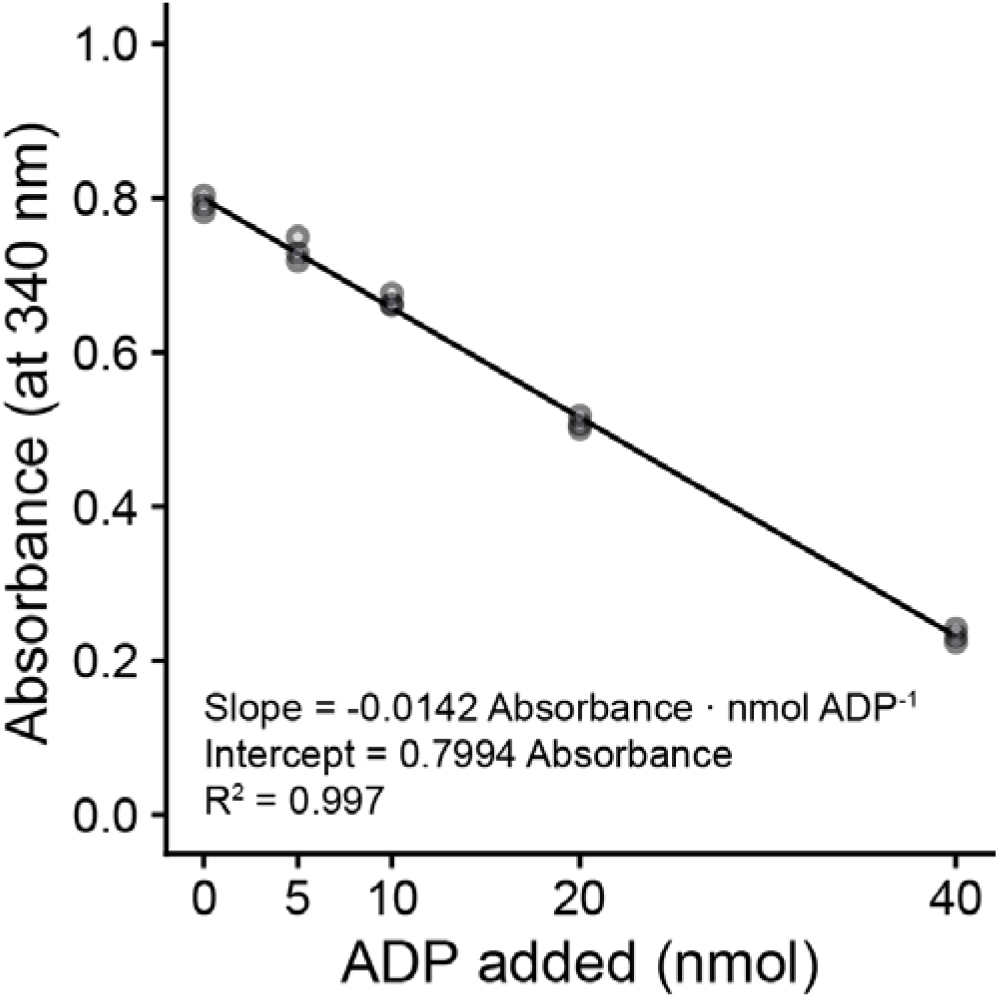
Example standard curve used to estimate activity of the Na^+^/K^+^-ATPase. Activity of the Na^+^/K^+^-ATPase was estimated by measuring depletion of NADH. To translate disappearance of NADH to the production of ADP production by ATPases, we created a standard curve correlating absorbance (i.e. NADH concentration) to predetermined amounts of ADP produced (or added in the case of the standard curve).

**Figure S2.**
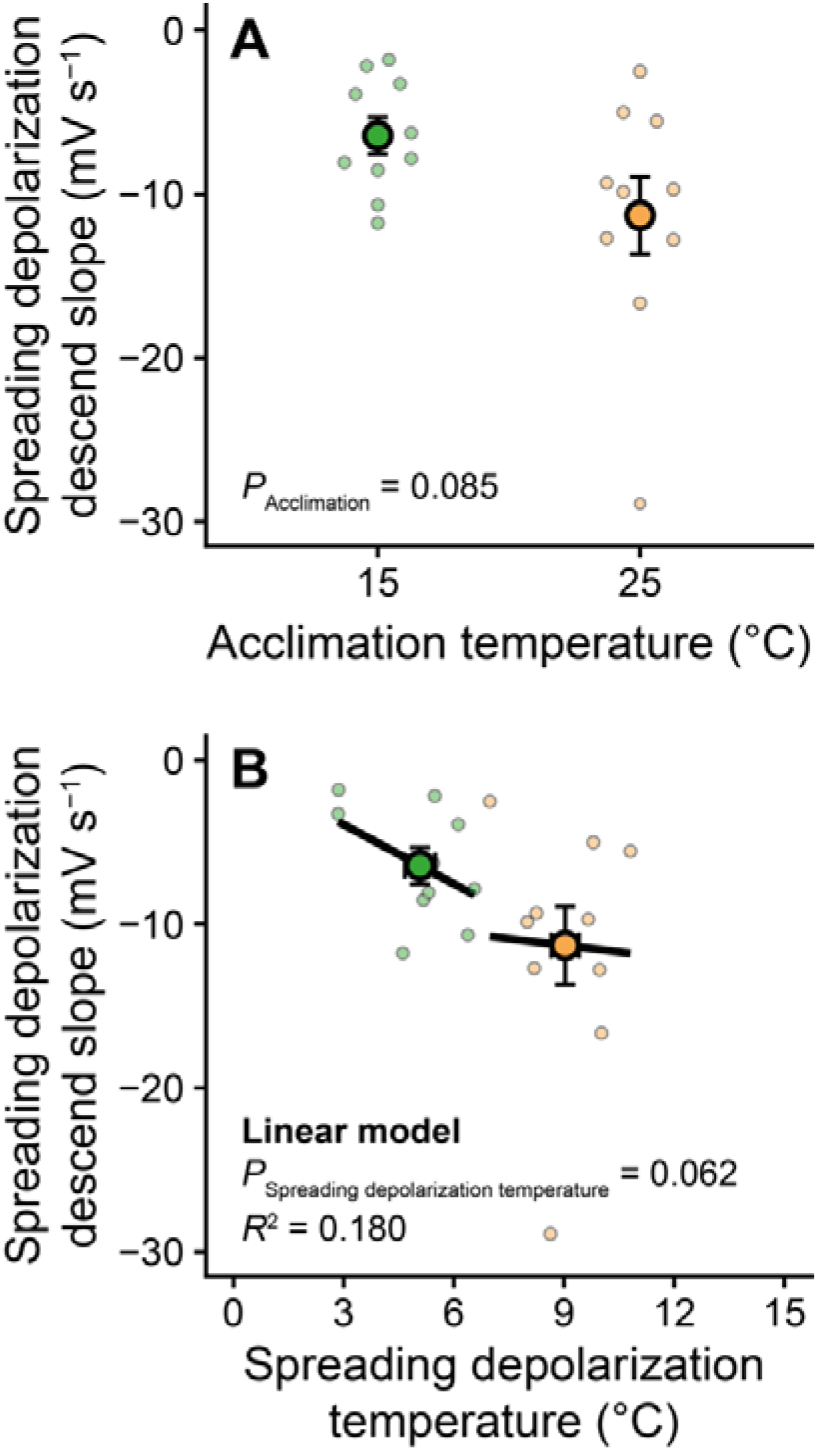
Cold acclimation reduces the slope of the drop in transperineurial potential during spreading depolarization. (A) Cold acclimation (15°C; green) of female *Drosophila melanogaster* reduces the magnitude of the spreading depolarization descend slope relative to warm acclimation (25°C; orange), despite not reaching the level of statistical significance. (B) A negative relationship between spreading depolarization temperature and descend slope of the transperineurial potential was evident; however, this too did not reach the level of statistical significance. N = 10 flies per group, which are depicted by smaller, translucent points, and group means are depicted as large opaque symbols.

**Figure S3.**
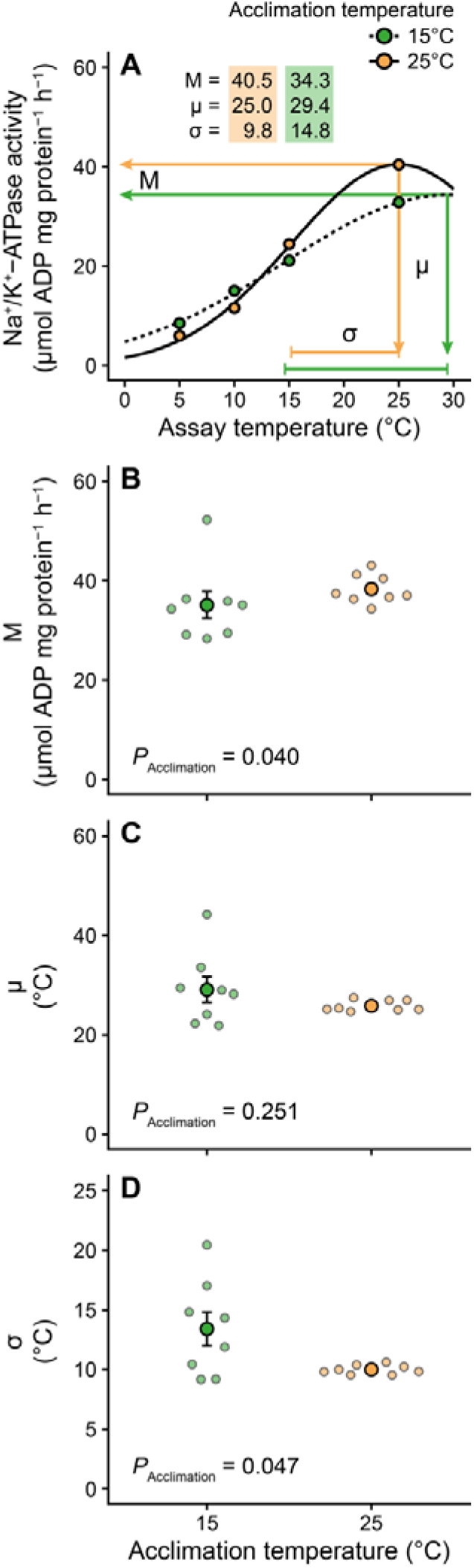
Three-parameter Gaussian model estimates for the temperature-dependent activity of the Na^+^/K^+^-ATPase. Because the relation between temperature and Na^+^/K^+^-ATPase activity was not exponential we decided to also fit a three-parameter Gaussian model to the data, to more accurately describe the trend. (A) A visual representation of each parameter in the model superimposed on data from two samples. This analysis was performed for each sample such that means could be compared between the two acclimation groups. The analysis revealed that (B) cold acclimation led to a lower maximum activity of the Na^+^/K^+^-ATPase at the optimal temperature, (C) the temperature for optimal Na^+^/K^+^-ATPase was unaffected by acclimation temperature, and (D) the activity of the Na^+^/K^+^-ATPase was more resilient to lowered temperature after cold acclimation (larger σ denotes that a bigger change in temperature is needed to reduce activity). Note that the Gaussian model often (10/16 samples) relied on extrapolation beyond the data (more often in the cold-acclimated group, which might explain the higher degree of variation), as is evident from example data from a cold-acclimated fly in panel A.

